# MultipleXed Population Selection and Enrichment single nucleus RNA sequencing (XPoSE-seq) enables sample identity retention during transcriptional profiling of rare populations

**DOI:** 10.1101/2023.09.27.559834

**Authors:** Katherine E. Savell, Rajtarun Madangopal, Padmashri Saravanan, Ryan G. Palaganas, Kareem D. Woods, Drake J. Thompson, Olivia R. Drake, Megan B. Brenner, Sophia J. Weber, Elise Van Leer, Jae H. Choi, Toni L. Martin, Jody C. Martin, Mia K. Steinberg, James W. Austin, Chloé I. Charendoff, Bruce T. Hope

**Affiliations:** Intramural Research Program, National Institute on Drug Abuse, Baltimore, MD 21224; BD Biosciences, San José, CA 95131

## Abstract

Single nucleus RNA-sequencing is critical in deciphering tissue heterogeneity and identifying rare populations. However, current high throughput techniques are not optimized for rare target populations and require tradeoffs in design due to feasibility. We provide a novel snRNA pipeline, Muliple**<u>X</u>**ed **<u>Po</u>**pulation **<u>S</u>**election and **<u>E</u>**nrichment snRNA-sequencing (**XPoSE-seq**), to enable targeted snRNA-seq experiments and in-depth transcriptomic characterization of rare target populations while retaining individual sample identity.

---

Single cell and nucleus RNA-sequencing (sc/sn-RNAseq) approaches disentangle the heterogeneity of complex tissues and aid in the identification of rare populations otherwise masked by conventional bulk-tissue procedures. While current pipelines effectively characterize common populations, there are multiple challenges to perform snRNA-seq on rare target populations^1^. High-throughput technologies isolate large nuclei numbers, but without enrichment, rare target populations are masked by majority non-target populations. When enrichment is used to increase target population frequency, multiple biological replicates are pooled in a single capture, sacrificing critical individual subject information. However, recent work highlights the importance of true biological replicates to properly account for between-replicate variation and reduce false discoveries in single cell/nucleus analyses^2^.To address these issues, we developed XPoSE-seq, which combines flow cytometry-based rare population enrichment with an antibody-based multiplexing snRNA-seq strategy to optimize cost, throughput, and sample identity retention (**Figure 1A**). The key component in XPoSE-seq is a novel reagent (XPoSE-tag) that leverages an antibody against the ubiquitous nucleoporin complex protein (Nup62), found in nuclei from all cell types, with dual conjugations of 1) a far-red dye (R718) to identify Nup62-labeled nuclei in flow cytometry and 2) one of eight distinct oligo-based Sample Tags (ST) to uniquely barcode nuclei from each sample. XPoSE-tag can be used in conjunction with other fluorescent labels to select/enrich similar proportions of target populations from each sample before mixing (multiplexing) samples prior to nuclei capture on the BD Rhapsody microwell-based system^3,4^.

**Figure 1:**
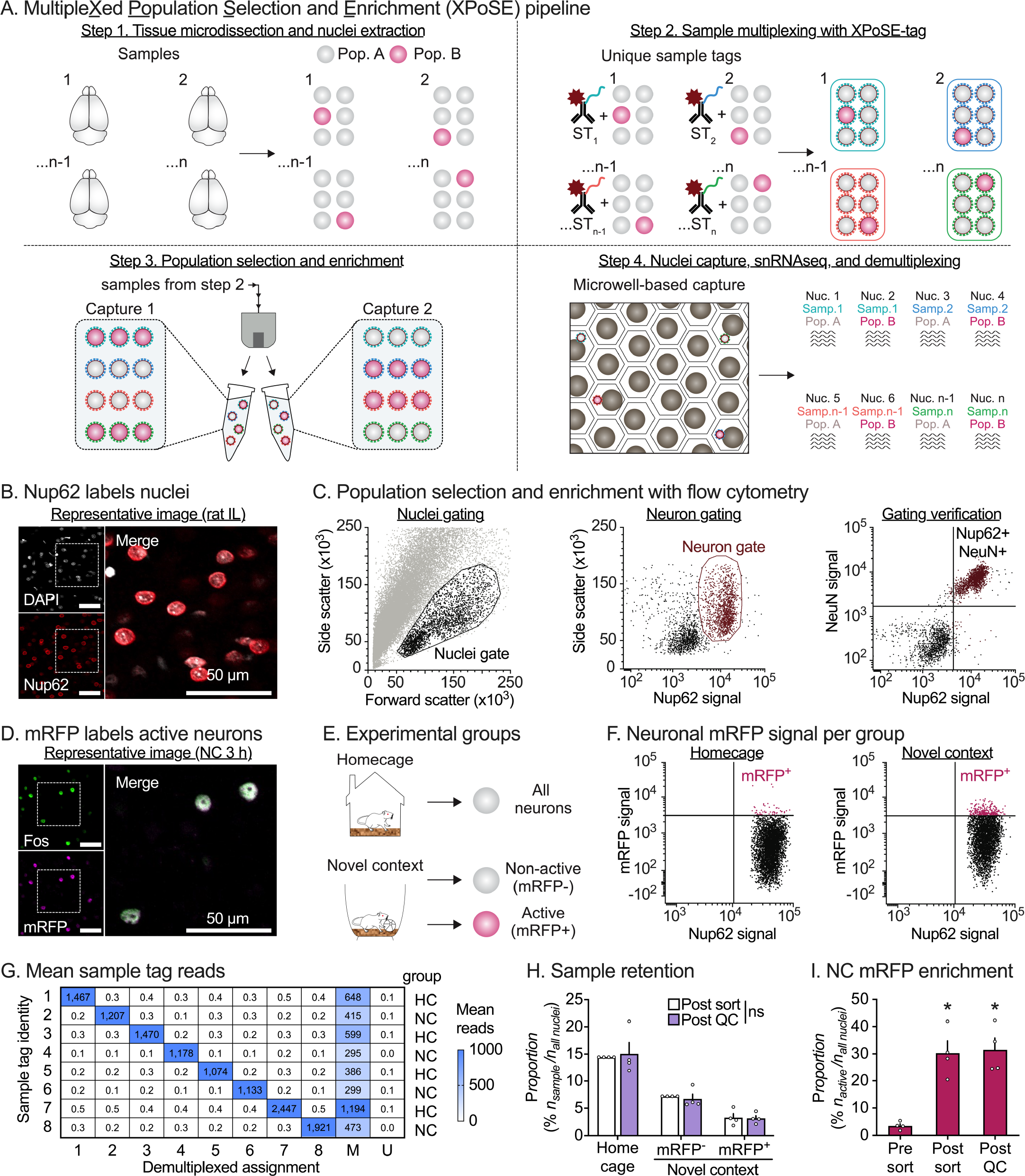
Development of Multiple<u>X</u>ed <u>Po</u>pulation <u>S</u>election and <u>E</u>nrichment single nucleus RNA-sequencing. **A**, XPoSE-seq workflow. After tissue microdissection and nuclei isolation, nuclei from individual samples are incubated with a R718 dye- and oligo-conjugated nucleoporin antibody (XPoSE-tag). Specific populations are enriched by gating for the XPoSE-tag and other population-labeling markers with flow cytometry. Sorted nuclei containing known proportions of each sample/population are captured in a cartridge-based system (BD Rhapsody). **B**, Representative immunofluorescent (IF) image of rat IL shows successful nuclei (DAPI-labeled, white) co-labeling with nucleoporin (red). Scale bar = 50 μm. **C**, Event scatter plots from nuclear preps demonstrate dual-conjugated Nup62 is detected with flow cytometry. Nuclei are gated from debris based on forward- and side-scatter properties (left). Neuronal populations are selected by nucleoporin signal (middle). Neurons comprise the right nucleoporin positive population as verified by co-staining for the neuronal nuclei marker, NeuN (right). **D**, Representative IF image of rat IL shows that mRFP is expressed in cells expressing Fos in *Fos-mRFP* transgenic rats. **E**, Experimental design (left), *Fos-mRFP* transgenic rats explored a novel context for one hour or remained in the homecage. All neurons were collected in Homecage samples, and Active (mRFP^+^) and Non-active (mRFP^-^) neuronal nuclei were collected from novel context samples. **F**, Gating strategy to enrich for behaviorally active neurons. The mRFP-positive threshold is determined by comparing mRFP signal between neurons from Novel context and Homecage preps. **G**, Demultiplexed assignment matches to presence of corresponding Sample tag identity reads. **H**, Sample proportion between sorted and Post-QC total nuclei was not affected by multiplexing, regardless of population size. Data is expressed as the mean percent ± S.E.M. of each rat/population combination in the dataset. ns indicates no difference (*p* > 0.9999) between the percent collected and retained from each rat (*n* = 4 per group). **I**, The mean percentage ± S.E.M. of mRFP in the neuron population is enriched from the original sample by sorting and is retained in the dataset after QC. * Significant difference from original sample (*n* = 4 per group). See Table S1 for detailed listing of all statistical outputs.

We verified that XPoSE-tag reliably labeled nuclei (identified using DAPI fluorescence) in the infralimbic (IL) subregion of rat medial prefrontal cortex (**Figure 1B**). Flow cytometry of IL nuclei preparations (**Figure 1C**, left) revealed two distinct Nup62-positive populations within the nuclei gate (**Figure 1C**, center). After co-staining for the neuronal nuclear protein NeuN, we found that larger-sized nuclei were NeuN-positive (i.e., neurons) while smaller-sized nuclei were NeuN-negative (non-neurons). This size difference between neurons and non-neurons agrees with previous studies^5^, and allows for XPoSE-tag based selection of either subpopulation via flow cytometry (**Figure 1C**, right).

As a proof of principle study, we applied XPoSE-seq to enrich and profile the transcriptome of behaviorally active, Fos-positive neurons, a rare population that typically makes up <5% of neurons in a brain region^6^. We labeled behaviorally active neurons using *Fos-mRFP* transgenic rats^7^, where expression of the fluorescent protein mRFP is temporally induced in neurons that express the endogenous activity marker Fos (**Figure 1D**, labeling timecourse **Figure S1**). Nuclei preps were prepared from samples taken 2 h after 1 h of novel context (NC) exploration (or from homecage controls). Using fluorescence-activated nuclei sorting (FANS), we sorted Active (mRFP-positive) and Non-active (mRFP-negative) neuronal nuclei from four rats in the novel context group (**Figure 1E-F**). We sorted neurons agnostic of mRFP signal from four homecage (HC) rats to serve as baseline control. We counterbalanced populations from the 8 rats between two nucleus captures, multiplexing 8 populations per capture using the BD Rhapsody system. After library preparation, sequencing, and alignment/demultiplexing, we confirmed robust XPoSE-tag labeling, with ST reads corresponding to expected sample assignment (**Figure 1G**). There were no differences in population retention across the three collected populations (**Figure 1H**), indicating that the pipeline can be used to enrich for and retain both abundant (Control or Non-active) and rare (Active) populations from individual subjects. While the objective was to collect a 1:1 ratio of Active to Non-active neurons in NC subjects, we were ultimately limited by the size of the Active population. Nevertheless, the proportion of Active neurons from NC subjects was enriched approximately 9-fold (**Figure 1I**) compared to the original sample (**Table S1** for statistical outputs).

Dimensionality reduction of sorted nuclei expression counts confirmed majority neuronal clusters and negligible glia contamination (1.7%). Cluster transcriptomes matched previously identified excitatory (Slc17a7-positive) and inhibitory (Gad1-positive) cortical neuron classes (**Figure 2A & Figure S2**, public web browser at https://nidairpneas.shinyapps.io/xpose) with excellent reproducibility in cell-type distribution between cartridge captures (**Figure S3**). We examined the cell-type distribution of each population (**Figure 2B**) and found that NC Active nuclei were enriched in *IT-L5/6* and *Sst* clusters and underrepresented in *CT-L6, NP-L5/6, Sst Chodl, Vip*, and *Lamp5* neuronal nuclei clusters (**Figure 2C**). As expected, there were no differences in cell-type proportions between abundant HC and NC Non-active nuclei samples (Statistical outputs in **Table S1**).

**Figure 2:**
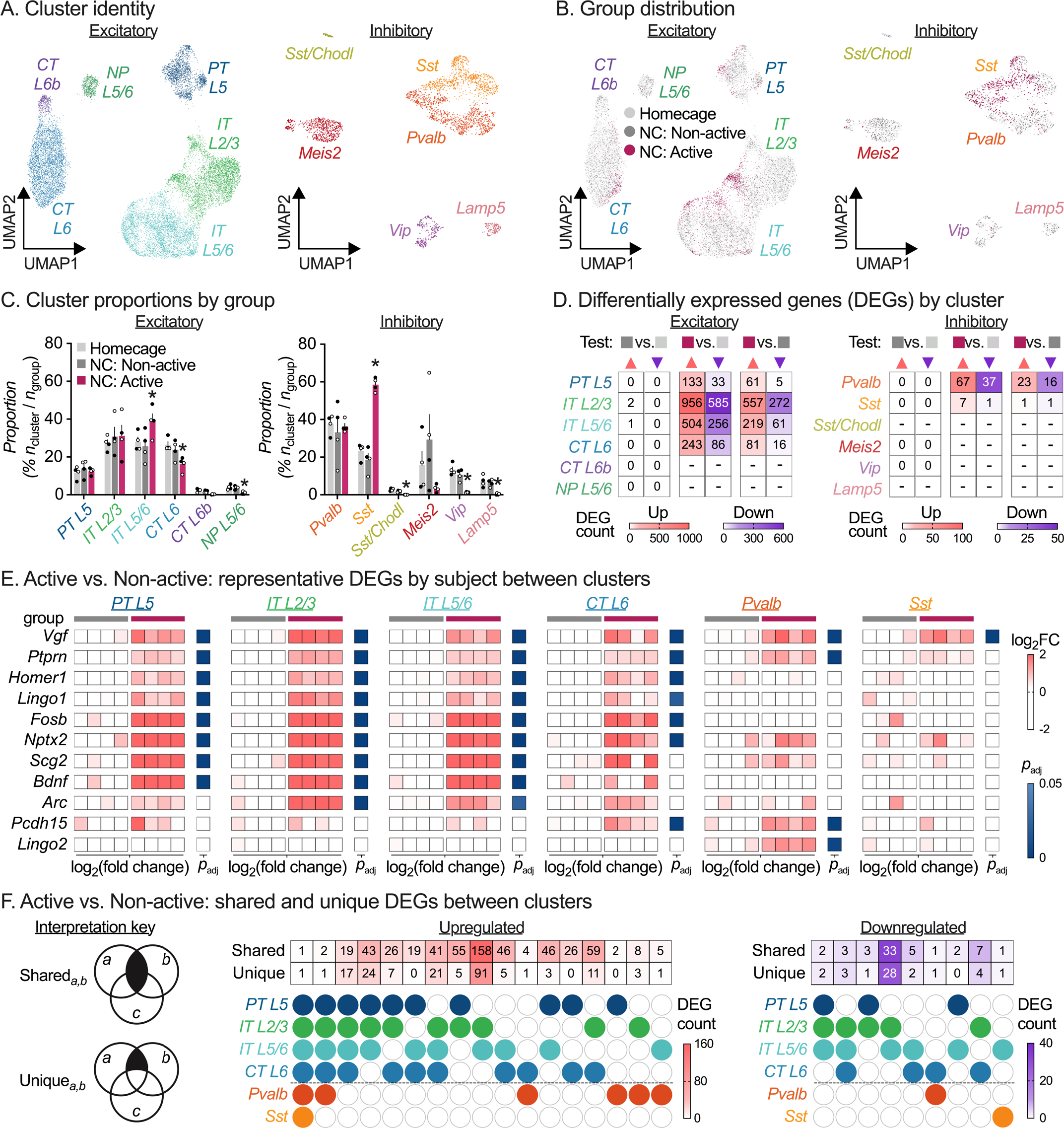
Application of XPoSE-seq to study behaviorally active neurons. **A**, UMAP representation of excitatory (left) and inhibitory (right) neuronal cell type clusters in the rat infralimbic cortex. UMAP, Uniform Manifold Approximation and Projection; PT, pyramidal tract; IT, intratelencephalic; CT, corticothalamic; L, layer. UMAP distribution (**B**) and mean percent ± S.E.M. (**C**) by group for excitatory and inhibitory clusters. Open circles indicate male samples, closed circles indicate female samples (*n* = 4 / group, 2 F, 2 M). *Significant differences (FDR-adjusted *p* < 0.05) multiple paired *t*-tests (Active vs. Non-active). **D**, Summary heatmap showing counts of differentially expressed genes (DEGs) per cluster (cluster inclusion criteria of 75 nuclei / group, FDR-adjusted < 0.05) for Non-active 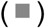 vs. Homecage 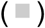, Active 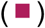 vs. Homecage 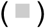, and Active 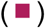 vs. Non-Active 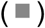 comparisons. Upregulated genes are in salmon 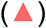, downregulated genes in purple 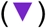. **E**, Representative gene examples of differentially expressed genes for Active 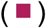 vs. Non-Active 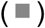 comparisons across clusters showing biological replicates. Data is shown for each biological replicate as log_2_FC of normalized counts, FDR-adjusted < 0.05. **F**, Gene set comparisons (Shared and Unique) by cell type for DEGs in the Active vs. Non-active comparison. See **Table S1** for detailed listing of all statistical outputs, and **Table S2** for cluster/comparison specific DEGs.

A key benefit of XPoSE-seq is that individual subject metadata is retained for each population, allowing differential expression (DE) analyses that account for biological replicates and associated variation. We performed cluster-specific DE analysis on pseudo-bulked counts from each biological replicate (*n* = 4 / group),excluding clusters with fewer than 75 nuclei / population. Transcriptional changes were found overwhelmingly in the Active population (**Figure 2D, Table S2**), with differentially expressed genes comprising many known activity-dependent genes (**Figure 2E**)^6,8-10^. Next, we investigated shared or unique transcriptional changes across cell types (**Figure 2F**). Surprisingly, while several genes showed enrichment across multiple active clusters, only a single gene, *Vgf*, was upregulated in all active clusters. Additionally, while *Sst* was overrepresented in the Active population, there were no other transcriptional changes within this cell type. Finally, in line with and extending previous bulk-sample based studies^9,11^, activity-dependent transcriptional changes were almost non-existent in the Non-active population compared to Homecage nuclei, irrespective of cell-type cluster.

In the past decade, single-cell technologies have illuminated diverse cell types and cell states within tissues across species, discovering novel and rare populations via high-throughput transcriptomic analyses. XPoSE-seq complements these efforts, pinpointing rare-population-specific transcriptomic signatures and merging them with sample-specific metadata like sex, treatment, or behavior. XPoSE-tag, an antibody-based^12^ multiplexing extension, captures rare populations from multiple samples in user-defined proportions. In this proof-of-concept study, XPoSE-seq effectively enriched and maintained sample-specificity for neurons activated in a novel context. Active neurons were distributed across IL cell types and had both shared and cell-type specific gene expression patterns. This underscores XPoSE-seq’s utility in exposing cell-type-specific alterations within rare populations.

Many studies opt to combine all biological replicates of an experimental group into a single capture step due to cost and feasibility. However, this approach risks overrepresentation bias in terms of collected cell clusters or differentially expressed genes (DEGs). Furthermore, it limits differential expression (DE) analysis, as many studies equate biological replicates with the number of cells/nuclei, inaccurately inflating analysis power. XPoSE-seq offers a solution by enabling individual subjects to serve as biological replicates, allowing precise assessment of each sample’s contribution to specific analyses. Moreover, it enables precise control over population proportions prior to nucleus capture, essential for mitigating unequal sample sizes and resulting issues like unequal variance and reduced statistical power. XPoSE-seq is well positioned to support targeted multi-sample transcriptomic profiling of rare populations labelled using transgenic reporter lines, cell-type and circuit specific viral tools, or emerging enhancer-driven virus-based approaches^13,14^.

In summary, XPoSE-seq combines the benefits of antibody-based sample-multiplexing and FACS-based population enrichment with high-throughput snRNAseq technologies to enable cost-effective transcriptomic characterization of rare target populations while retaining sample identity information. In addition, it provides fine control over proportions of nuclei collected per population and sample and allows for more robust statistical testing approaches. The XPoSE-seq approach and experimental strategy employed here is broadly applicable to investigation of other target populations, such as precious samples, rare cell-types, altered cell states by treatment or disease, or combinations thereof.

## Acknowledgements

We thank members of Hope lab for their support and insight during all stages of this study. We thank Ueta lab (University of Occupational and Environmental Health, Kitakysushu, Japan) for providing the *Fos-mRFP* transgenic line. We thank Dr. Francois Vautier and the NIDA IRP Transgenic Breeding staff for management of the *Fos-mRFP* transgenic lines, and we thank Dr. Christopher Richie (NIDA IRP Genetic Engineering and Viral Vector Core) and Madeline Merriman for confirming genotype of *Fos-mRFP* transgenic rat sublines. We thank the NIH Intramural Sequencing Core for assistance with sequencing. We thank Dr. BaDoi Phan for generating the mm10 annotation liftoff to Rn7.2 used during alignment.

## Author contributions

K.E.S., R.M. and B.T.H. designed the experiments. K.E.S., and R.M. conceptualized the XPoSE-seq approach. J.C.M. generated XPoSE-tag reagent. K.E.S. and R.G.P. validated the XPoSE-tag reagent. R.M., R.G.P., and O.R.D., and M.B.B., ran behavioral experiments and collected samples for XPoSE-seq. K.E.S., R.G.P., R.M., C.I.C., and J.W.A., ran XPoSE-seq experiments. K.E.S., R.M., S.J.W., E.V.L., J.H.C., and R.G.P. performed Fos-mRFP rat validation experiments. K.E.S., P.S., M.K.S., R.M., K.D.W., D.J.T., T.L.M., and R.G.P., established analysis pipelines and analyzed the data. P.S., D.J.T., and K.E.S. developed R Shiny web application. K.E.S., R.M., and B.T.H. wrote the paper. All authors reviewed and approved the final version prior to submission.

## Data, Materials, and Code Availability

All relevant data that support the findings of this study are available by request from the corresponding author (B.T.H.). Sequencing data will be deposited in Gene Expression Omnibus at time of publication. Custom R analysis scripts to reproduce the figures are available on the XPoSE Github repository (https://github.com/ksavell/XPoSE). A public web browser to explore the dataset is located at https://nidairpneas.shinyapps.io/xpose.

## Competing Interests

J.C.M., M.K.S., J.W.A, and C.I.C. were employees of BD Biosciences. Manuscript approval by BD Biosciences was not required, and BD Biosciences had no influence regarding data analysis, data interpretation, and discussion. All other authors declare that they do not have any conflicts of interest (financial or otherwise) related to the text of the paper. This work was supported by the Intramural Research Program of the National Institute on Drug Abuse. K.E.S and R.M. received funding from the NIH Center for Compulsive Behaviors. K.D.W., and O.R.D were supported by the NIDA IRP Scientific Director’s Fellowship for Diversity in Research. P.S., D.T., E.V.L, and J.H.C. were supported by the NIH Summer Research Program, and T.L.M was supported by the NIDA Undergraduate Research Internship Program.

## Open Access

This article is licensed under a Creative Commons Attribution 4.0 International License, which permits use, sharing, adaptation, distribution, and reproduction in any medium or format, as long as you give appropriate credit to the original author(s) and the source, provide a link to the Creative Commons license, and indicate if changes were made. The images or other third-party material in this article are included in the article’s Creative Commons license, unless indicated otherwise in a credit line to the material. If material is not included in the article’s Creative Commons license and your intended use is not permitted by statutory regulation or exceeds the permitted use, you will need to obtain permission directly from the copyright holder. To view a copy of this license, visit http://creativecommons.org/licenses/by/4.0/.

© The Author(s) 2023

## Methods

### Subjects

We used male and female *Fos-mRFP*^*+/-*^ transgenic rats on Wistar background (NIDA transgenic breeding facility, n = 54), weighing 250-450 g in all experiments. We group-housed rats (two to three per cage) in the animal facility and single-housed them before the experiments. For all experiments, we maintained the rats under a reverse 12:12 h light/dark cycle (lights off at 8 AM) with free access to standard laboratory chow and water in their home cages, throughout the experiment. All procedures were approved by the NIDA IRP Animal Care and Use Committee and followed the guidelines outlined in the Guide for the Care and Use of Laboratory Animals^1^. We report the number of rats included in each experiment in the corresponding figure legend.

### Behavioral procedures

We used novel context exposure for acute neuronal activation based on previous studies where this procedure has been shown to induce robust IL activation^2^. We also collected brains directly from the homecage to serve as baseline controls. The procedures for each experiment are outlined below.

#### Fos-mRFP labeling timecourse

We exposed rats to a novel environment for 1 h, and then placed them back in the homecage for varying periods of time before perfusions and brain extraction. We randomly assigned rats to 6 groups and collected PFA perfused brain tissue at 1, 2, 3, 4, 8, or 24 hours after the start of novel context exposure. We also collected brains from a separate group of rats directly from the homecage to serve as baseline controls. During this experiment, we discovered a copy number discrepancy between breeding sublines, and 16 subjects were excluded from the experiment through copy number determination with ddPCR.

#### XPoSE pipeline demonstration

We exposed rats to a novel environment for 1 h, and then placed them back in the homecage for 2 h to allow for mRFP expression (peak expression at 3 h). We used isoflurane to induce a light plane of anesthesia prior to rapid decapitation and fresh brain tissue extraction. We also collected brains from a separate group of rats directly from the homecage to serve as baseline controls.

### Sample collection for immunohistochemistry (Fos-mRFP timecourse, XPoSE-tag validation)

We anesthetized rats with isoflurane (at varying time periods after behavioral testing, or from homecage) and perfused them transcardially with ∼250 ml of 1x PBS at pH 7.4 (PBS), followed by ∼250 ml of 4% paraformaldehyde at pH 7.4 (PFA). We extracted brains and post-fixed for an additional 1-4 h in PFA before transferring them to 30% sucrose for 48 h at 4 °C. We froze sucrose equilibrated brains on dry ice and stored them at −80 °C prior to sectioning.

### Immunohistochemistry (Fos-mRFP timecourse, XPoSE-tag validation)

We used a cryostat (Leica, Model CM3050 S) to collect coronal sections (40 μm) containing infralimbic cortex into PBS, and stored them at 4 °C until further processing. We rinsed free-floating sections first in PBS with 0.5% Tween20 and 10 μg/ml heparin (wash buffer, 3 × 10 min), incubated them in PBS with 0.5% TritonX-100, 20% DMSO, and 23 mg/ml glycine for 3 h at 37 °C (permeabilization buffer), and then in PBS with 0.5% TritonX-100, 10% DMSO, and 6% normal donkey serum (NDS) for 3 h at 37 °C (blocking buffer) prior to antibody labeling. We diluted primary (1°) antibodies (Fos: anti-Phospho-c-Fos (Ser32) (D82C12) XP® Rabbit mAb, #5348, Cell Signaling Technology, RRID: AB_10557109, 1:2500; mRFP: anti-DSRed mouse mAb, sc-390909, Santa Cruz Biotechnology, RRID: AB_2801575, 1:1000; Nucleoporin: anti-Nucleoporin 62 Mouse mAb, 610497, BD Biosciences, RRID: AB_397863, 1:500) in PBS with 0.5% Tween20, 5% DMSO, 3% NDS, and 10 μg/ml heparin (1° Ab buffer) and incubated sections in this solution overnight at 37 °C. Following, 1° antibody labeling, we rinsed sections in wash buffer (3 × 10 min) and then incubated in secondary (2°) antibodies (Fos: Alexa Fluor® 647 Donkey Anti-Rabbit IgG (H+L), Jackson ImmunoResearch Labs, 711-605-152, RRID: AB_2492288, 1:250; mRFP: Alexa Fluor® 488 Donkey Anti-Mouse IgG (H+L), Jackson ImmunoResearch Labs, 715-545-150, RRID: AB_2340846, 1:250; Nucleoporin: Alexa Fluor® 647 Donkey Anti-Mouse IgG (H+L), Jackson ImmunoResearch Labs, 715-605-150, RRID: AB_2340862, 1:250) diluted in PBS with 0.5% Tween20, 3% NDS, and 10 μg/ml heparin overnight at 37 °C. Following, 2° antibody labeling, we rinsed sections again in wash buffer (3 × 10 min), mounted them onto gelatin-coated slides, allowed them to partially dry. We then coverslipped the slides using MOWIOL or Vectashield Vibrance mounting medium with DAPI nuclear stain and allowed them to hard-set overnight prior to imaging on a confocal microscope (Nikon C2, or Olympus FLUOVIEW FV3000).

### Confocal imaging (Nucleoporin validation)

We acquired confocal images of infralimbic cortex (AP + 3.0 to AP +3.2, relative to Bregma) on an Olympus FLUOVIEW FV3000 confocal microscope using a 20x/0.75 NA air objective and 2x area zoom. The imaged field of view was 318.2 × 318.2 μm^2^ (3.2181 pixels/μm) with 8.0-μs dwell time and 140 μm pinhole. For DAPI fluorescence, we excited tissue with 0.9% laser power at 405 nm and collected emitted fluorescence from 430 to 470 nm with a detection PMT voltage of 530. For Nup62 fluorescence, we excited tissue with 0.5% laser power at 640 nm and collected emitted fluorescence from 650 to 750 nm with a detection PMT voltage of 500. We imaged the two channels sequentially (order: 405, then 640) to minimize overlap and acquired 4 images (2 hemispheres x 2 sections) per animal.

### Confocal imaging (Fos-mRFP timecourse)

We acquired confocal images of infralimbic cortex (AP + 3.0 to AP +3.2, relative to Bregma) on a Nikon C2 confocal microscope using a 20x/0.75 NA air objective. The imaged field of view was 642.62 × 642.62 μm^2^ (1.5935 pixels/μm) with a 5.3-μs dwell time and 30 μm pinhole. For mRFP fluorescence, we excited tissue with 2% laser power at 488 nm and collected emitted fluorescence from 500 to 550 nm with a detection gain of 80 and an offset of −1. For Fos fluorescence, we excited tissue with 2% laser power at 640 nm and collected emitted fluorescence from 670 to 1000 nm with a detection gain of 105 and an offset of 5. We imaged the two channels sequentially (order: 488, then 647) to minimize overlap and acquired 4 images (2 hemispheres x 2 sections) per animal for analysis.

### Sample collection and storage for XPoSE pipeline

We snap froze fresh brains in pre-chilled isopentane at -80 °C for 15 seconds, wrapped the frozen brain samples in labeled aluminum foil wrappers, and submerged them under dry ice for short term storage. We used a -80 °C freezer for long term frozen brain sample storage.

### IL microdissection

We used a cryostat (Leica CM 3050S) to section the brains at -20 °C. First, we mounted frozen brains (−80 °C) onto cryostat pedestals and allowed them to equilibrate to -20 °C prior to sectioning. We cut 300 μm coronal sections containing IL (AP +2.8 to AP +3.8), micro dissected IL tissue from both hemispheres, and stored tissue pieces in nuclease free 1.5 mL polypropylene microcentrifuge tubes (Eppendorf) at -80 °C until further processing.

### Nuclei isolation and XPoSE-tag labeling

We used established detergent-mechanical cell lysis protocols^3^ for single nuclei dissociation from frozen tissue punches with some modifications as described below. Briefly, we transferred frozen tissue punches to a prechilled Dounce homogenizer tube and added ice cold detergent lysis buffer (0.32 M sucrose, 10 mM HEPES pH 8.0, 5 mM CaCl_2_, 3 mM MgAc, 0.1 mM ETDA, 1 mM DTT, 0.1% Triton X-100). We lysed tissue and released nuclei (10 strokes of pestle A, then 10 strokes of pestle B). We diluted the lysate in chilled low sucrose buffer (0.32 M sucrose, 10 mM HEPES [pH 8.0], 5 mM CaCl2, 3 mM MgAc, 0.1 mM ETDA, 1 mM DTT) then filtered the suspension through a 40 μm strainer to remove debris, centrifuged the sample at 3,200 x g for 10 min at 4 °C, and poured off the supernatant. Next, we resuspended the pellet from each sample in 750 mL of chilled resuspension buffer (1X PBS, 0.4 mg/mL BSA, 0.2 U/μL RNAse Inhibitor [Biosearch Technologies, Cat. 30281]) containing one of 8 unique XPoSE-tags (Nucleoporin 62 antibody conjugated to R718 fluorescent dye and one of eight distinct oligo-based Sample Tags (ST) to uniquely barcode nuclei from each sample, custom reagent, BD Biosciences, 1:2000). For verifying the neuronal nuclei gate, we added both the XPoSE-tag and neuronal nuclear marker NeuN (anti-NeuN Mouse mAb clone A60, MAB377X, Millipore Sigma, RRID: AB_2149209, 1:500) We incubated the samples with antibody for 15 minutes at 4 °C on a rotating mixer, transferred to a 5 mL polystyrene round-bottom tube (Falcon) containing 3.5 mL chilled resuspension buffer, centrifuged at 250 × g for 10 min at 4 °C, and aspirated 4 mL supernatant containing residual XPoSE-tag. We resuspended the pellet containing XPoSE-tag labelled nuclei in 1-2 mL chilled resuspension buffer and passed through a 40 μm filter prior to fluorescence activated nuclei sorting (FANS).

### Fluorescence activated nuclei sorting (FANS)

We used FANS to isolate XPoSE-tag labelled (Nup62+) neuronal nuclei, and to enrich for Nup62+/mRFP+ nuclei in our samples. We used a BD FACS Melody sorter equipped with 3 excitation lasers (488 nm, 561 nm and 640 nm) and 9 emission filters for FANS. We employed a sequential gating strategy similar to previous studies (Hope lab). Nuclei formed a distinct cluster that separated from debris and was gated in forward scatter area vs. side scatter area view. For XPoSE-tag (Nup62-BDR718) detection, we excited samples with 640 nm laser and collected emitted fluorescence from 690 to 750 nm with a detection PMT voltage of 605, gating the larger Nup62 positive population (neurons). For native mRFP detection, we excited samples with 561 nm laser and collected emitted fluorescence from 595 to 631 nm with a detection PMT voltage of 603. We used mRFP rats from the *Homecage* group to define a threshold gate for *active* (mRFP+) nuclei. We used these gates to sort *active* (mRFP+) and *non-active* (mRFP-) neuronal nuclei from each *Novel context* group rat into separate 1.5 mL Protein Lo-Bind Eppendorf tubes containing 750 μL of chilled resuspension buffer (sorting block chilled with recirculating chiller throughout sorting). To calculate mRFP percentages, we recorded 2,000 events from each sample before initializing the sort. We collected on average 3400 *active* (mRFP+) and 7500 *inactive* (mRFP-) neuronal nuclei per rat in the novel context group (n = 4) and counterbalanced the collected population between tubes so all experimental groups were represented in each capture. We collected 7500 neuronal nuclei per tube from each *Homecage* group rat (n = 4) to generate the IL cell-type atlas and serve as baseline controls. We processed all 8 rats on the same day and collected nuclei into the same 2 × 1.5 mL Eppendorf tubes - thus each tube contained up to 7500 neuronal nuclei (mRFP+, mRFP-, or all neuronal nuclei, counterbalanced) from each rat at the end of nuclei collection and was processed using a BD Rhapsody separate cartridge.

### Nuclei capture and barcoding

We used BD Rhapsody Single cell analysis platform for nuclei capture and followed manufacturer’s suggested protocols for nuclei staining, cartridge preparation, sample loading, bead loading, lysis and reverse transcription steps. We first added Vibrant Dye Cycle Green nuclear dye (Thermo Fisher, Catalog number V35004, 1:1000) to each Eppendorf tube (containing collected nuclei) and incubated the samples for 10 minutes on ice. We diluted samples to 1.5 ml with BD Sample Buffer with RNAse Inhibitor, then centrifuged the tubes at 250 × g for 10 min at 4 °C. We removed excess buffer to a final volume of ∼650 μL. We used two cartridges for the experiment - one for each 1.5 mL Eppendorf tube used during sorting. We primed the BD cartridge with absolute ethanol, and wash buffers and then loaded resuspended nuclei. We incubated the loaded cartridge for 30 minutes at 4 °C and then used the BD Rhapsody Scanner to scan the cartridge and estimate nuclei capture efficiency. Next, we loaded barcoded capture beads onto the cartridge and incubated beads with the sample as directed by manufacturer protocol. We then performed washes to remove excess beads and ran a second cartridge scan to determine bead load efficiency. After the scan, we engaged the bottom magnet in ‘Lysis’ mode to immobilize capture beads, added lysis buffer, and incubated for 2 min. We then switched the magnet to ‘Retrieval’ mode, followed manufacture protocols to collect beads with captured polyadenylated targets, and performed reverse transcription and an exonuclease reaction using the version 1 3’ kit (BD Biosciences, cat. 633733, 633773).

### Single nucleus RNA sequencing, demultiplexing and genome alignment

We followed manufacturer protocols to generate single-cell whole transcriptome mRNA, and Sample Tag libraries (BD Biosciences, cat. 633801) for sequencing on Illumina® sequencers. We followed manufacturer protocols to measure library concentration with Qubit 4 fluorometer (ThermoFisher, cat. Q33230) and length distribution with a Bioanalyzer 2100 (Agilent, cat. 5067-4626). We processed both cartridges in parallel and generated separate libraries that were indexed and pooled for sequencing. We sequenced whole transcriptome libraries at 45,000 reads per nucleus and sample tag libraries at 1,000 reads per nucleus on an Illumina Novaseq S4 lane (PE 150 bp). We trimmed sequences to 75 bp then used the BD Rhapsody WTA Analysis Pipeline (v1.11 rev 8, Single-Cell Multiplex Kit - Mouse) on Seven Bridges Genomics platform to demultiplex raw sequencing reads and align to the reference rat genome (Rn7.2). We included introns and exons during alignment and generated filtered count matrices with recursive substitution error correction (RSEC) based on a liftover from mm10 mouse genome annotation to Rn7.2 rat genome^4^. Sample tag UMIs and sample tag calls were generated for each nuclei ID.

### snRNAseq analysis

All analysis scripts to reproduce the figures are on our XPoSE repository on Github (https://github.com/ksavell/XPoSE). Rhapsody count matrices with RSEC for each cartridge were analyzed with Seurat (v4.3) in R (v4.3.0). Seurat objects were created from count matrices, and the sample tag ID and sample tag counts were assigned as metadata from the corresponding sample tag outputs. Experimental metadata for each nucleus was assigned based on the cartridge/XPoSE-tag combination, and cells with multiple XPoSE sample tags or fewer than 50 gene features were removed.

Of the 21,871 nuclei that passed QC filtering, we normalized and scaled the data using the 2000 most highly variable features before applying dimension reduction. We screened the cells for known markers for neurons and non-neuronal populations (*Snap25* [pan-neuronal], *Slc17a7* [excitatory neuron], *Gad1* [inhibitory neuron], *Mbp* [oligodendrocyte], *Gja1* [astrocyte], *Col5a3* [microglia]) and removed 553 (1.7% of the total) nuclei as non-neuronal contamination. We then split excitatory and inhibitory neuron populations into two objects to process in parallel. After subsetting, we reclustered both excitatory and inhibitory data and generated the top 10 marker genes for each cluster using FindMarkers function in Seurat to visualize as a heatmap in **Supplemental Figure 1**. We manually annotated the clusters based on marker expression using the Allen Cell Type Database nomenclature^5^.

For differential expression analysis, Libra (v1.0)^6^ was used to sum gene counts for all nuclei for each rat per population to create a pseudobulked gene expression matrix for each cluster with > 75 nuclei per group in the comparison. Any genes with counts < 5 were removed for analysis. DESeq2 (v1.40) ^7^ was used to calculate DEGs for each cluster and comparison using false discovery rate corrected p-values. All DEGs for each comparison are listed in **Supplemental Table 2**. Normalized counts were extracted from the DESeq2 object to calculate individual subject fold change for representative genes. ComplexUpSet was used to create a combination matrix (using both distinct and intersect modes) of DEG list comparisons. While we included both female and male subjects in accordance with sex as a biological variable policy, the current experiment is not appropriately powered to find differential gene expression by sex.

### Statistical Analysis

Statistical and graphical analysis were performed using R and GraphPad Prism (version 9.5.1). We tested the data for sphericity and homogeneity of variance when appropriate. When the sphericity assumption was not met, we adjusted the degrees of freedom using the Greenhouse–Geisser correction. Sample retention proportions were tested with 2-way repeated measures (RM) ANOVA with Bonferonni post-hoc. mRFP percentages/enrichment were tested using RM 1-way ANOVA with Tukey post-hoc. Proportions for excitatory and inhibitory clusters were tested using multiple paired *t-*tests in a *between* (Non-active vs. Homecage or Positive vs. Homecage) or *within* (Active vs. Non-active) subject design and corrected for multiple comparisons (FDR < 0.05) using p.adjust() in R. The mRFP labeling timecourse was tested using a 1-way ANOVA with Bonferonni post-hoc. Because our analyses yielded multiple main effects and interactions, we report only those that are critical for data interpretation. See **Table S1** for detailed listing of all statistical outputs.

## Supplementary Figures

**Figure S1.**
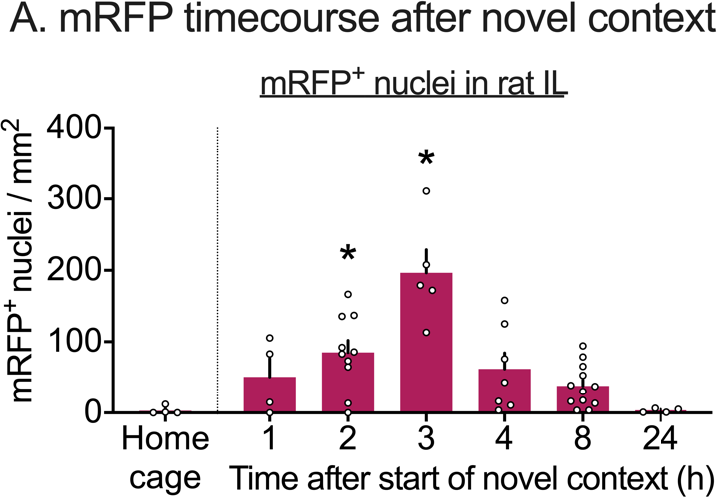
mRFP labels Fos-expressing active neurons in *Fos-mRFP* rats. **A**, Timecourse of mRFP labeling following 1 hour of novel context exploration in *Fos-mRFP* rats (*n* = 4 – 12 per group). Data is expressed as means ± S.E.M. * significant difference from homecage (HC). See Table S1 for detailed listing of all statistical outputs.

**Figure S2.**
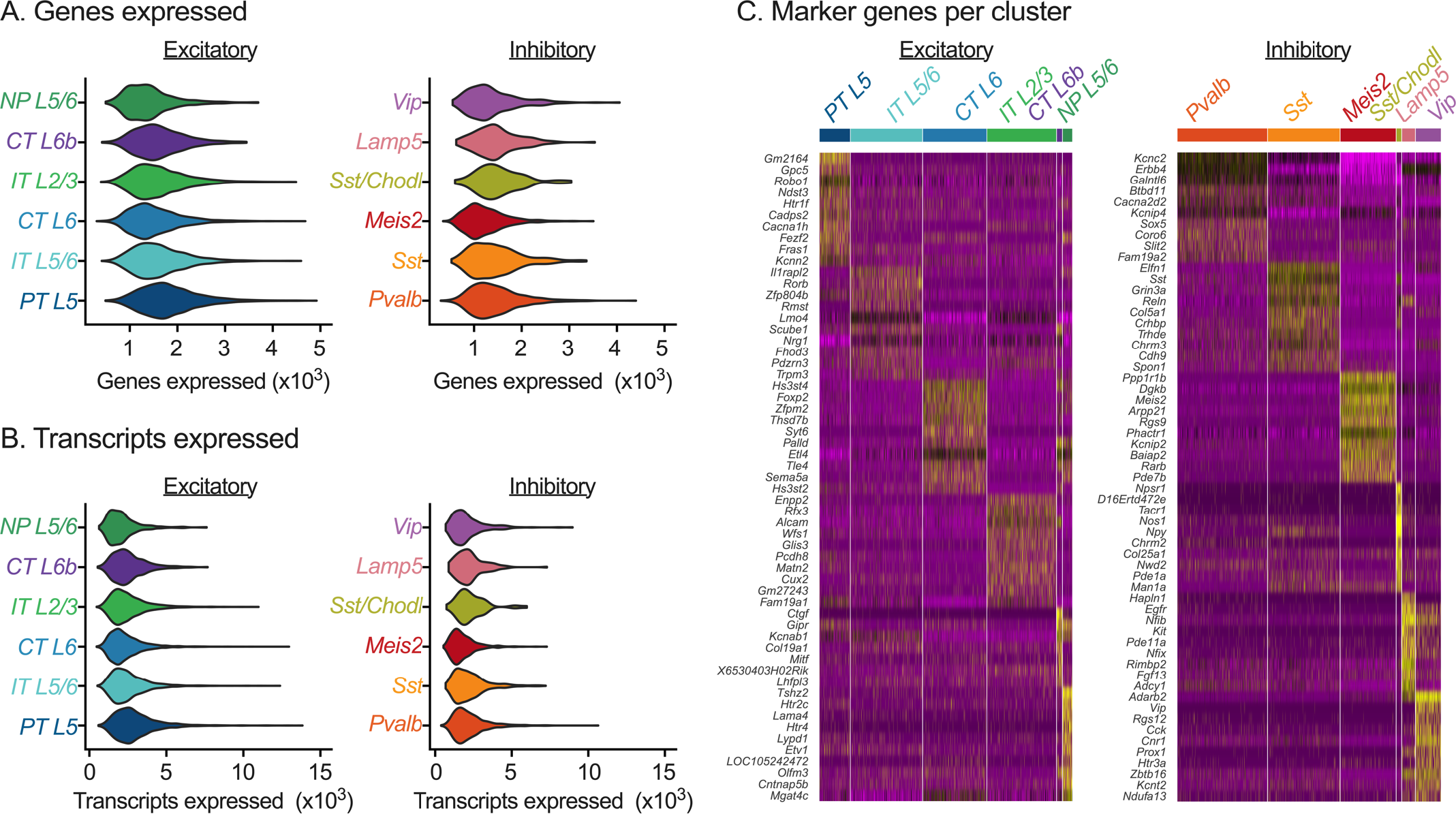
Quality control and cluster markers of excitatory and inhibitory neuron clusters. (**A**) Genes expressed in excitatory and inhibitory neuron clusters. Violin plot shows the number of genes expressed in excitatory (left) and inhibitory (right) cluster types collapsed across biological replicates and populations (n = 8). (**B**) Total number of transcripts expressed. Violin plot shows the total number of transcripts expressed in excitatory (left) and inhibitory (right) cluster types. (**C**) Heatmap of top ten genes expressed in major excitatory (left) and inhibitory (right) neuron clusters of the rat IL.

**Figure S3.**
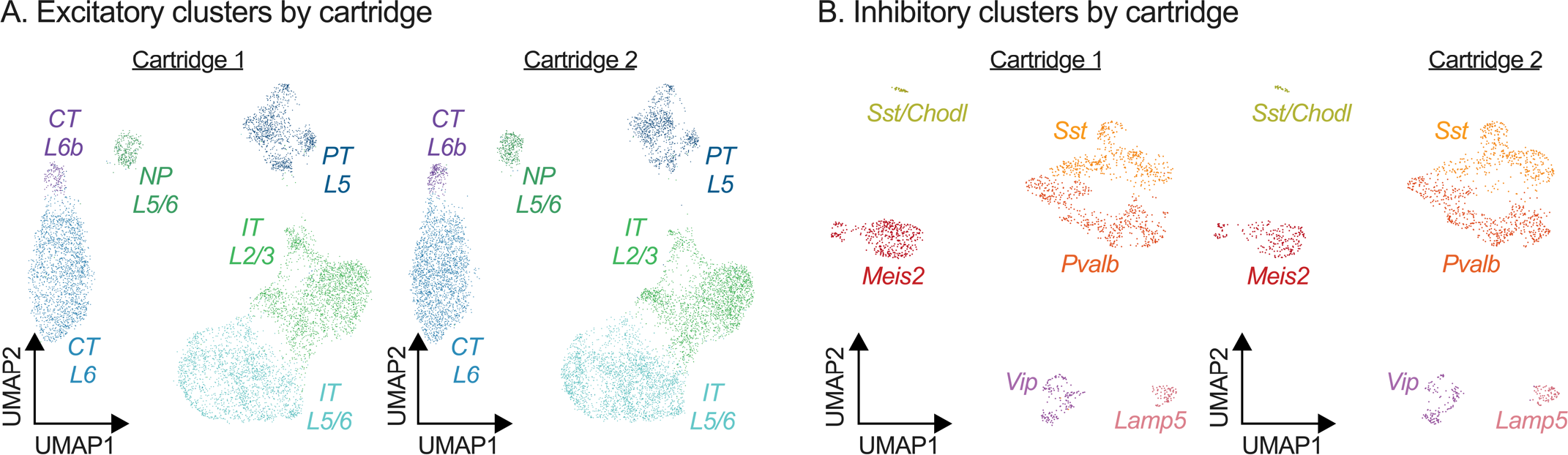
Neuronal cell type cluster type distribution between cartridges. (**A**) UMAP representation of excitatory neuron cluster distribution by cartridge. (**B**) UMAP representation of inhibitory neuron clusters by cartridge.

## Notes

https://nidairpneas.shinyapps.io/xpose

https://github.com/ksavell/XPoSE

